# Vertical sleeve gastrectomy lowers kidney SGLT2 expression in the mouse

**DOI:** 10.1101/741330

**Authors:** Elina Akalestou, Livia Lopez-Noriega, Isabelle Leclerc, Guy A. Rutter

**Affiliations:** Section of Cell Biology and Functional Genomics and Imperial Diabetes Network, Division of Diabetes, Endocrinology and Metabolism, Imperial Centre for Translational and Experimental Medicine, Department of Medicine, Imperial College London, London W12 0NN, United Kingdom

## Abstract

**Background:** Bariatric surgery has been established to improve insulin sensitivity and glucose clearance, but also increases insulin and glucagon secretion. Each of the above effects have also been observed following treatment with sodium glucose co-transporter 2 (SGLT2) inhibitors.

**Aim:** To determine whether there is an effect of bariatric surgery (Vertical Sleeve Gastrectomy; VSG) on renal SGLT2 expression in mice.

**Methods:** Eighteen lean mice underwent VSG (n=8) or sham (n=9) surgery. Glucose tolerance tests with or without treatment with the SGLT2 inhibitor dapagliflozin were performed four weeks post operatively, in order to assess if pharmacological SGLT2 inhibition has the same euglycemic effects after bariatric surgery. Kidneys were harvested from fed mice and SGLT2 expression was analysed using Quantitative reverse-transcription PCR and immunofluorescence.

**Results:** VSG mice displayed significantly improved glucose tolerance (AUC=103±6.8; AUC=66.6±2.9 in control and VSG mice, respectively; p<0.001), despite an absence of significant weight loss when compared to sham operated mice (p=0.37, Mann-Whitney test). Treatment of sham-operated mice with dapagliflozin (10 mg/kg) improved glucose tolerance. In contrast, dapagliflozin did not further improve glucose tolerance in VSG-operated mice. Moreover, qRT-PCR and immunofluorescence analysis on mouse kidneys demonstrated a significant lowering of SGLT2 expression at both the mRNA (n=7, p<0.0001) and protein (n=5, p=0.0007) levels four weeks after VSG.

**Conclusions:** Vertical sleeve gastrectomy in lean animals causes a significant inhibition of SGLT2 expression in the kidney cortex. These findings are in line with our previous results on the effects of Duodenal Jejunal Bypass in lean rats, and point towards a physiologically-relevant gut-kidney axis. SGLT2 inhibition may thus be an important mechanism through which bariatric surgery improves glucose tolerance in man.

## Introduction

Type 2 Diabetes (T2D) is characterised by decreased β-cell mass and function, leading to defective insulin secretion in the face of lowered hormone sensitivity ^1^. Despite an abundance of pharmacological and nutritional breakthroughs in recent years ^2-6^, there is still no permanent cure to T2D. Within the last 15 years, bariatric surgery, which was originally conceived as an intervention for weight loss^7-9^, was also remarkably effective in reversing T2D^10, 11^. Thus, since 2007, a number of randomised clinical trials have reported that bariatric surgery results in health benefits beyond weight loss and adipose tissue reduction, including improvements in T2D, steatohepatitis, cardiovascular health, cancer and fertility ^12-15^ and therefore pointing towards a direct or indirect trans-organ communication with the gastrointestinal tract. Numerous studies have attempted to shed light on these lines of communication and consequently mechanisms of disease remission, in the hope of replicating bariatric surgery with less invasive, pharmacological treatments.

One of the mechanisms hypothesised to contribute to the T2D remission following surgery is the change in intestinal glucose absorption. Studies investigating the effect of Roux-en-Y Gastric Bypass (RYGB) and Vertical Sleeve Gastrectomy (VSG) reported different mechanism between the two surgeries, with glucose trapped within the intestinal epithelial cells of rats and humans that underwent RYGB and reduced intestinal absorption of alimentary glucose following VSG ^16^. Moreover, in humans, glucose absorption was found only to occur in the common limb of the RYGB, while overall postprandial glucose concentration was reduced post operatively ^17, 18^. Of note, there have been contradicting results with regards to intestinal sodium glucose transport protein 1 (SGLT1) expression levels ^19-22^, which is the primary glucose transporter responsible for intestinal glucose absorption, along with glucose transporter-2 (GLUT2/SLC2A2).

A recently released class of anti-diabetic drugs focusses on the reduction of glucose absorption by inhibiting the renal glucose transporter, known as sodium-glucose transport protein 2 (SGLT2) ^23, 24^. Such SGLT2 inhibitors, or gliflozins, have proved to be hugely beneficial in patients with T2D ^25, 26^. They have also been observed to improve cardiovascular health, increase glucagon secretion and glycosuria ^27-29^. Interestingly, bariatric surgery has also been shown to improve T2D, cardiovascular health and increase glucagon ^30-32^. It was previously reported by one of us ^33^ that SGLT2 expression is down regulated in the kidney cortex following duodenal jejunal bypass in lean rats. In the present study, we aimed to assess the effect of VSG on SGLT2 expression in the mouse, a more genetically controllable species, and to investigate the existence of a kidney-gut axis in this species.

## Methods

### Animals

All animal procedures undertaken were approved by the British Home Office under the UK Animal (Scientific Procedures) Act 1986 with approval from the local ethical committee (Animal Welfare and Ethics Review Board, AWERB), at the Central Biological Services (CBS) unit at the Hammersmith Campus of Imperial College London. Adult male C57BL/6 mice (Envigo, Huntingdon U.K.) weighing 22-27g were maintained under controlled temperature (21-23°C) and light (12:12 hr light-dark schedule, lights on at 0700). The animals were fed PMI Nutrition International Certified Rodent Chow No. 5CR4 (Research Diet, New Brunswick, NJ) ad libitum. Animals were exposed to liquid diet (20% dextrose) three days prior to surgery and remained on this diet for up to seven days post operatively. All mice were divided in two groups, VSG (n=9) and sham (n=9), and were euthanized and harvested four weeks after surgery. Kidneys were harvested from all mice at four weeks following sham or VSG surgery in the fed state, and were either snap frozen in -80°C, fixed in formalin or both.

During the semaglutide study, adult male C57BL/6 mice were treated with either a single subcutaneous (SC) injection of semaglutide (Novo Nordisk UK) at 5nmol/ kg (n=6) or saline (n=6), for 7 days. Body weight was measured daily and all animals were euthanized and harvested on day 7.

### Vertical Sleeve Gastrectomy

Anaesthesia was induced and maintained with isoflurane (1.5-2%). A laparotomy incision was made and the stomach was isolated outside the abdominal cavity. A simple continuous pattern of suture extending through the gastric wall and along both gastric walls was placed to ensure the main blood vessels were contained. Approximately 60% of the stomach was removed, leaving a tubular remnant. The edges of the stomach were inverted and closed by placing two serosa only sutures, using Lembert pattern ^34^. The initial full thickness suture was subsequently removed. Sham surgeries were performed by isolating the stomach and performing a 1mm gastrotomy on the gastric wall of the fundus. All animals received a five-day course of SC antibiotic injections (Ciprofloxacin 0.1mg/kg).

### Glucose Tolerance Tests

Mice were fasted overnight (total 16 h) and given free access to water. At 0800, dapagliflozin (Forxiga, Astrazeneca, UK) was administered to the mice via oral gavage (10mg/kg suspended in 0.5% methylcellulose). At 1100, glucose (3 g/kg body weight) was administered via intraperitoneal injection. Blood was sampled from the tail vein at 0, 5, 15, 30, 60, 90 and 120 min after glucose administration. Blood glucose was measured with an automatic glucometer (Accuchek; Roche, Burgess Hill, UK).

### Plasma insulin and adiponectin measurement

To quantify circulating insulin and adiponectin levels, 100μl of blood was collected from the tail vein into heparin-coated tubes (Sarstedt, Beaumont Leys, UK). Plasma was separated by sedimentation at 10,000 g for 10 min (4°C). Plasma insulin levels were measured in 5μl aliquots by Crystal Chem (Zaandam, Netherlands) and adiponectin levels were measured by Abcam (USA).

### Immunohistochemistry of kidney sections

Isolated kidneys were fixed in 10% (vol/vol) buffered formalin and embedded in paraffin wax within 24 h of removal. Slides (5 μm) were submerged sequentially in Histoclear (Sigma, UK) followed by washing in decreasing concentrations of ethanol to remove paraffin wax. Permeabilised kidney slices were blotted with anti-rabbit SGLT2 (Abcam, USA) primary antibody. Slides were visualised by subsequent incubation with Alexa Fluor 488-labelled donkey anti-rabbit antibody. Samples were mounted on glass slides using VectashieldTM (Vector Laboratories, USA) and hard set with DAPI. Images were captured on a Zeiss Acio Observer.Z1 Motorised inverted widefield microscope fitted with a Hamamatsu Flash 4.0 Camera using a Plan-Apochromat 206/0.8 M27 air objective with Colibri.2 LED illumination. Data acquisition was controlled by Zeiss Zen Blue 2012 Software. Fluorescent quantification was achieved using Image J. Whole pancreas or kidney area was used to quantitate cell mass.

### RNA extraction, cDNA synthesis and Quantitative Polymerase Chain Reaction

Tissues were harvested and snap-frozen immediately in liquid nitrogen. RNA was purified using PureLink RNA kit (Thermo Fisher Scientific, UK). The purified RNA was dissolved in RNase and DNase free distilled water (Thermo Fisher Scientific, UK) and was immediately stored at -80°C until further analysis. Complementary DNA was synthesized from total RNA with High-Capacity cDNA Reverse Transcription Kit (Thermo Fisher Scientific, UK) according to the protocol recommended by the manufacturer. Quantitative Reverse Transcription PCR (qRT-PCR) analysis was used to quantify the expression level of SGLT2 and HNF1α in kidney cortex, and adiponectin in SC adipose tissue. Primers, which crossed a splice junction, were designed using Primer Express (Invitrogen, UK; Table 1). The expression levels were measured using Fast SYBR Green Master Mix (Invitrogen) and a 7500 Fast Real-Time PCR System (Applied Biosystems, UK) and calculated using the delta Ct method.

**Table 1:**
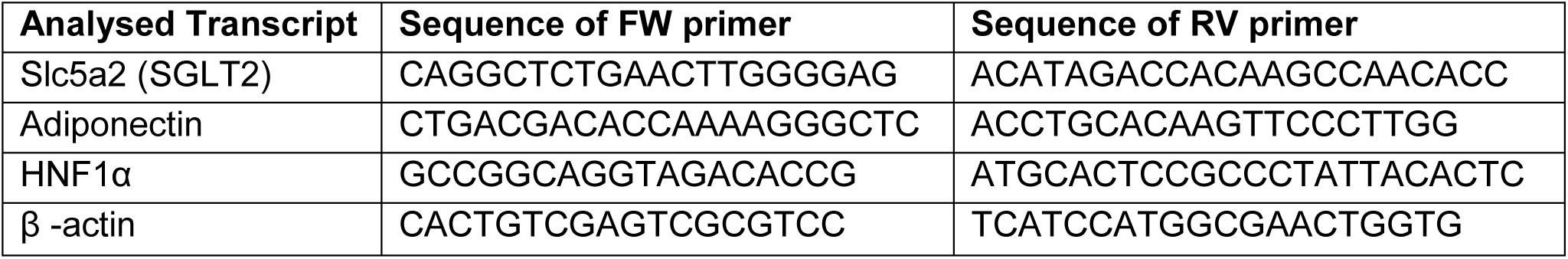
Primers Used for the Quantitative Detection of SGLT2, adiponectin, HNF1α, normalised to β-actin.

### Statistical Analysis

Data were analysed using GraphPad PRISM 7.0 software. Significance was tested using unpaired Student’s two-tailed t-tests with Bonferroni post-tests for multiple comparisons, or two-way ANOVA as indicated. P<0.05 was considered significant and errors signify ± SEM.

## Results

### Vertical Sleeve Gastrectomy improves glucose tolerance and insulin secretion without causing weight loss

Vertical Sleeve Gastrectomy significantly improved glucose tolerance in lean mice as assessed by intraperitoneal glucose tolerance test (IPGTT) four weeks post operatively (Fig 1A; AUC VSG 66.6 ±2.9; AUC sham 103 ±6.8,p<0.001 n=5-9). Remarkably, in VSG-treated mice, glucose peaked at 15 min. post glucose injection (3g/kg) and dropped to baseline levels within 60 min., whereas in sham operated mice glucose peaked at 30 min. and did not fully recover within the first 2 hours of measurement. This difference was associated with increased insulin secretion in VSG mice (Fig 1B) as the observed peak at 15 min. was almost three-fold higher when compared to sham mice (p<0.05). There was no significant difference of body weight between the two groups, as measured on the day of the experiment (Fig 1C).

**Figure 1.**
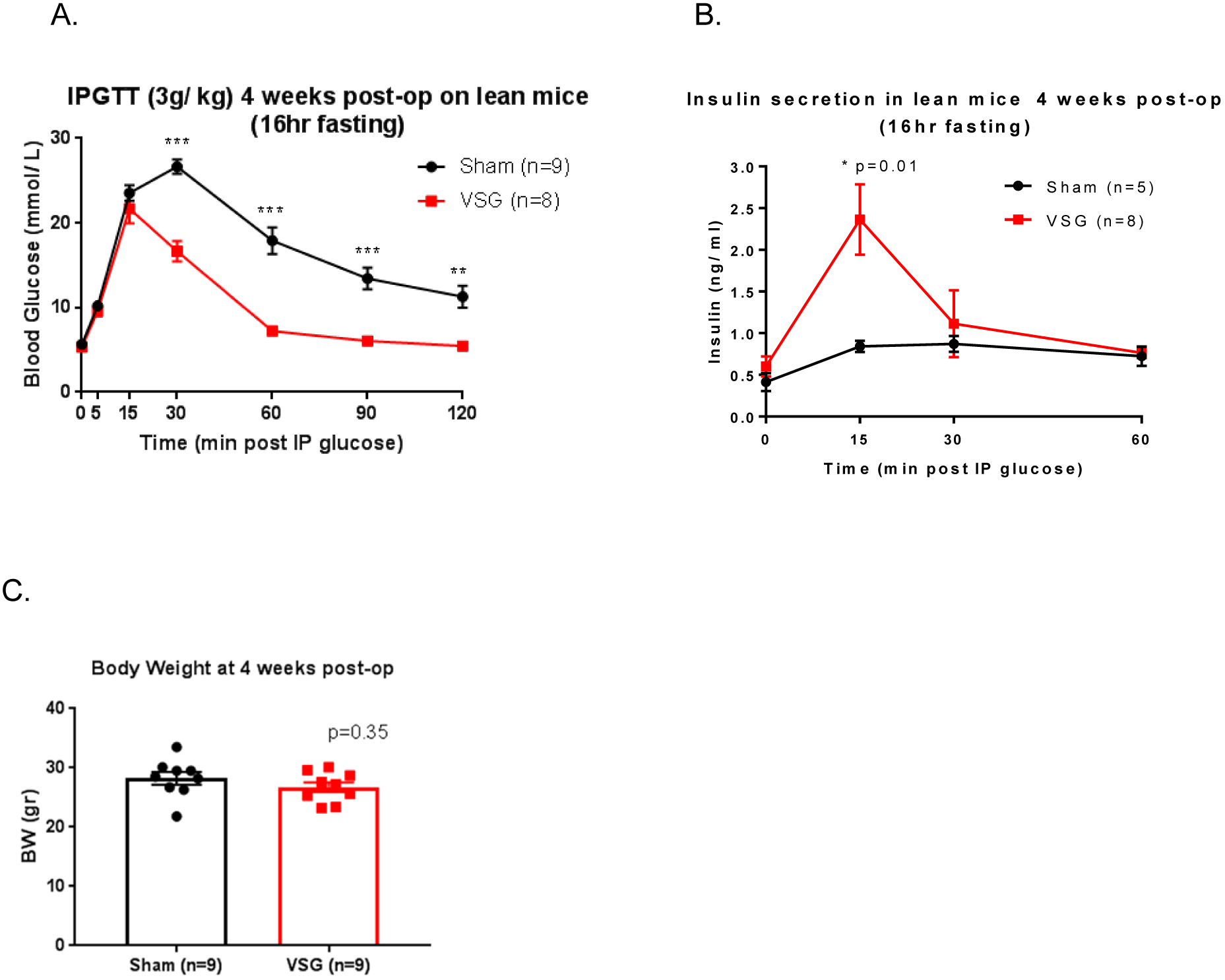
VSG in lean mice improves glucose tolerance and insulin secretion without changes in body weight. (A) Glucose was administered using intraperitoneal injection (3 g/kg) after mice were fasted overnight and blood glucose levels measured at 0, 5, 15, 30, 60, 90 and 120 min post injection, 4 weeks after surgery, n = 5–9 mice/group. (B) Corresponding insulin secretion levels measured on plasma samples obtained during the IPGTT performed in A (C) Body weight measurement of mice 4 weeks post surgery *P<0.05, ***P<0.001, by 2-way ANOVA. Data are expressed as means ± SEM.

### SGLT2 levels are lowered significantly in the kidney cortex following VSG in mice

When the kidney cortex was dissected and SGLT2 gene expression was measured using qRT-PCR, we observed a highly significant inhibition (p<0.001) of the expression of the *Slc5a2* gene following VSG (Fig 2A). This finding was confirmed at the protein level through immunohistochemistry revealing a 1.2-fold decrease after VSG (Fig 2B, C, D).

**Figure 2.**
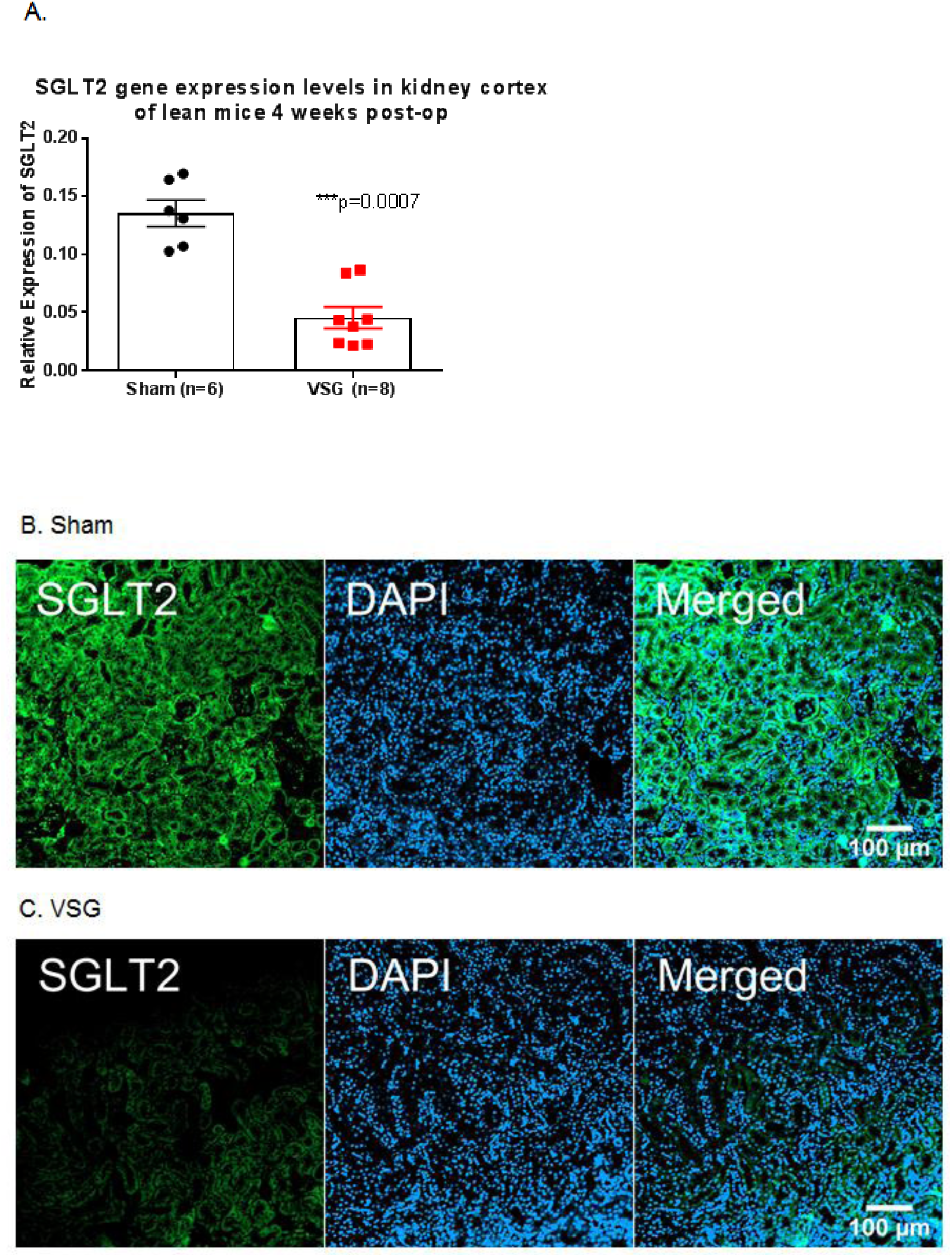
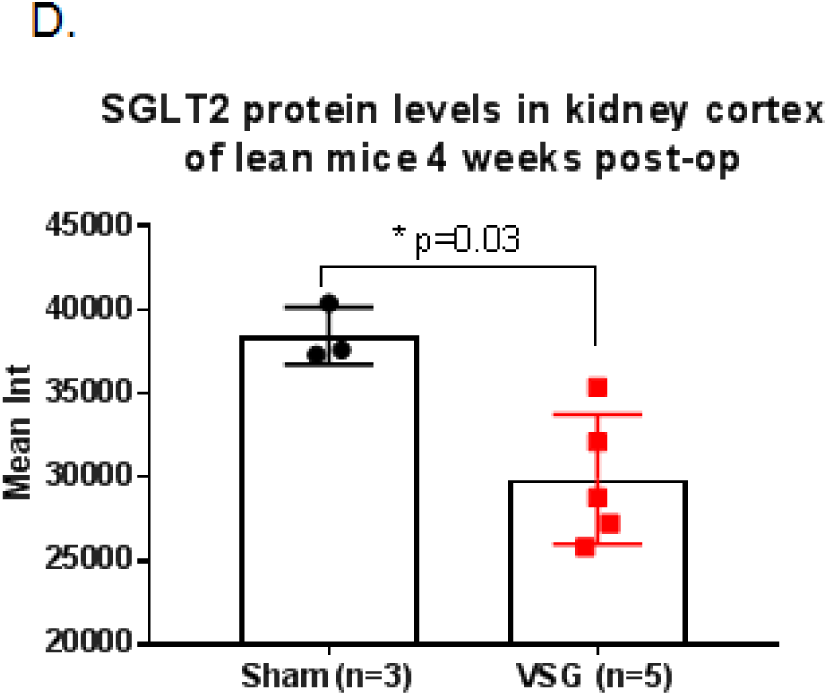
SGLT2 levels are decreased in the kidney cortex following VSG in mice (A) Quantitative PCR levels of SGLT2 gene expression in kidney cortex (B, C) Immunofluorescence staining of sham and VSG mice kidneys using anti-rabbit SGLT2 antibody (1:200; green) scale: 100um (D) Average Intensity measurement from immunofluorescence staining in kidney cortex. Each point represents the mean of 20-30 frames per slide, 3-4 slides per mouse. *P<0.05, by Student’s t-test; ***P<0.001 by 2-way ANOVA. Data are expressed as mean ns ± SEM

### Treatment with dapagliflozin has no additive effect on VSG to improve glucose tolerance

In order to further explore the reduction in SGLT2 expression observed in the VSG-treated animals, mice were administered with the SGLT2 inhibitor dapagliflozin (10mg/kg) via oral gavage three hours prior to an IPGTT (3g glucose/kg). In both sham and sham plus dapagliflozin groups, the glucose peak was observed at 30 min. post glucose injection. However, this peak was significantly lower in the Dapa group (Figure 3). Of note, peak glucose levels in the sham plus dapagliflozin group were significantly higher than the VSG group (p<0.001), suggesting that additional mechanisms contribute to the euglycaemic effect observed post-VSG. Nevertheless the glycemic excursions observed in the VSG and VSG plus dapagliflozin groups were almost identical, which indicates that treatment with dapagliflozin offers no additive effect to glucose lowering following bariatric surgery.

**Figure 3.**
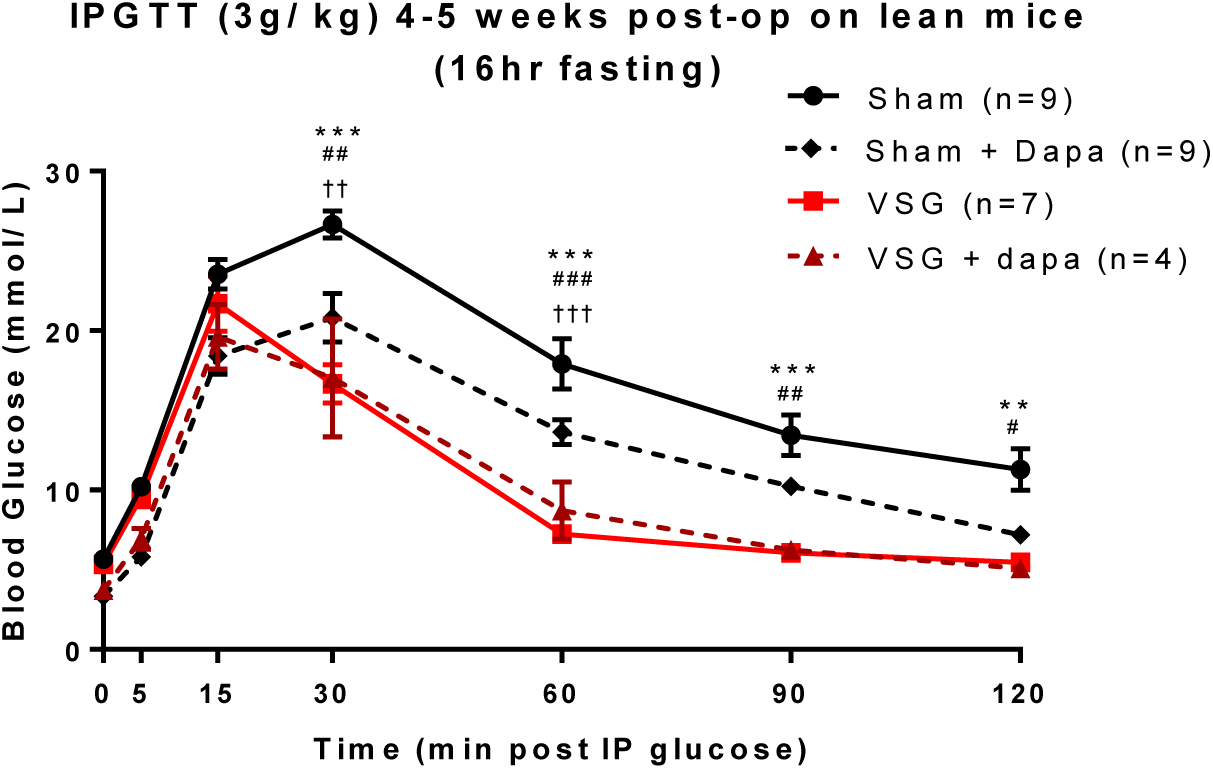
Dapagliflozin (10mg/ kg) was administered via oral gavage at -3hr after mice were fasted overnight, 4 weeks after surgery. Glucose was administered using intraperitoneal injection (3 g/kg) at time 0. Glycaemia was measured at 0, 5, 15, 30, 60, 90 and 120 min after glucose administration. 2-way ANOVA, * sham vs. VSG, # sham vs. VSG plus Dapa, † Sham plus Dapa vs VSG, **P<0.001, ***p<0.001 n = 4–9 mice/group.

### Semaglutide-induced reduced glycaemia does not lower SGLT2 levels significantly

Lean mice were treated daily with a SC dose (5nmol/kg) of the GLP-1 receptor agonist semaglutide for seven days so as to understand if the observed inhibited SGLT2 expression post bariatric surgery may be due to the lower basal glycaemia. The body weight of the mice was monitored daily and fed glycaemia was measured on day 7. Although body weight remained stable, glycaemia was significantly lower, when compared to the saline control group (Figure 4A, B). Nonetheless, kidney cortex SGLT2 gene expression was not significantly inhibited, albeit lower than the saline control (Figure 4C).

**Figure 4.**
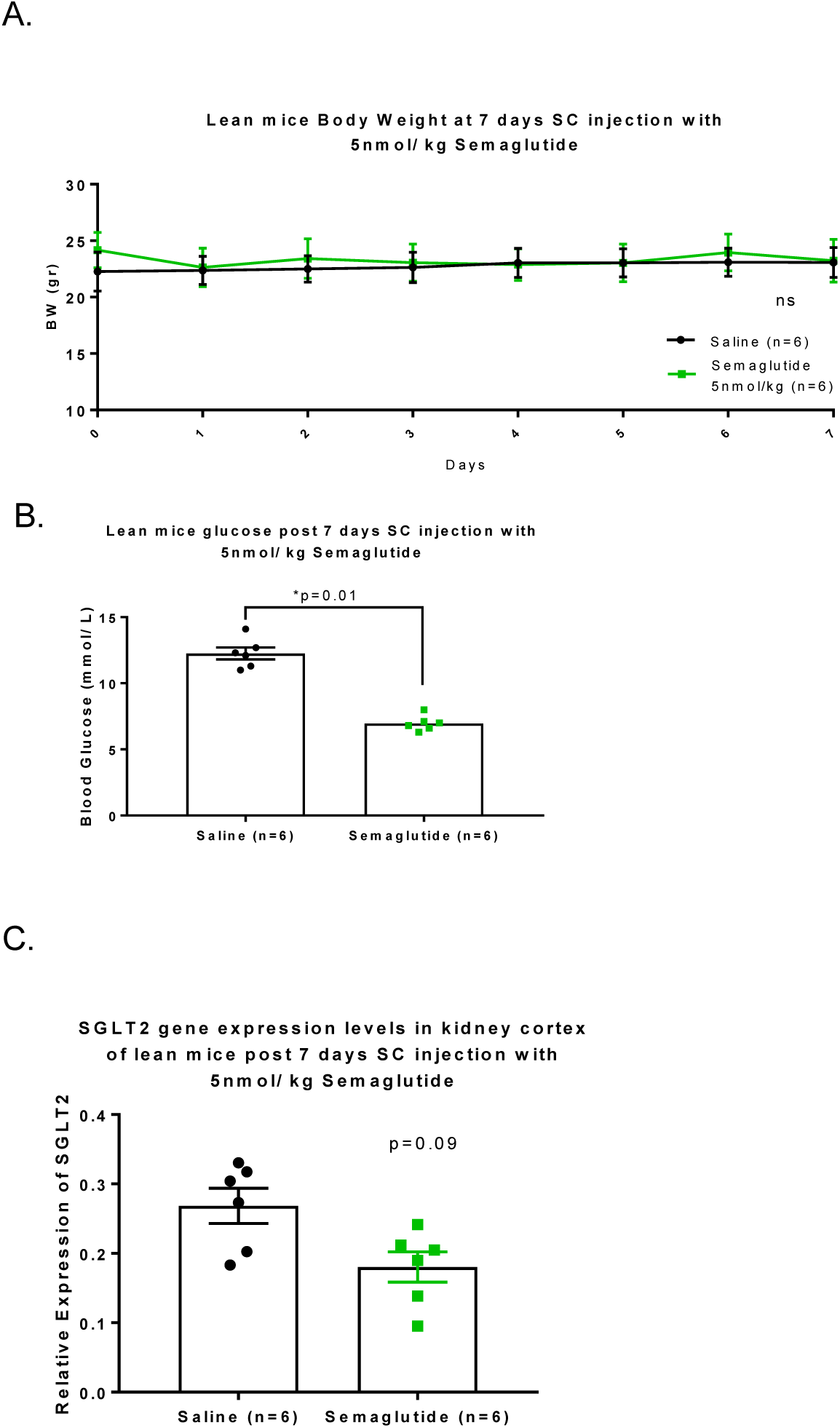
Semaglutide-induced glycaemia lowering does not decrease SGLT2 levels significantly. (A) Body weight measurement of mice that were treated with SC injection of 5nmol/kg semaglutide or saline for 7 days (B) fed glycaemia levels of mice on day 7 (C) Quantitative PCR levels of SGLT2 gene expression in kidney cortex. *P<0.05, by Student t-test. Data are expressed as means ± SEM

### VSG does not increase circulating adiponectin levels

Circulating adiponectin was measured in the plasma of fasted mice following bariatric surgery, as it has been previously reported that increased adiponectin reduces renal SGLT2 through activation of peroxisome proliferator-activated receptor delta (PPARδ) ^35^. However, adiponectin protein levels in plasma (Figure 5A) and adiponectin gene expression (Figure 5B) in SC adipose tissue were not significantly increased post-VSG, pointing towards alternative mechanisms of SGLT2 regulation.

**Figure 5.**
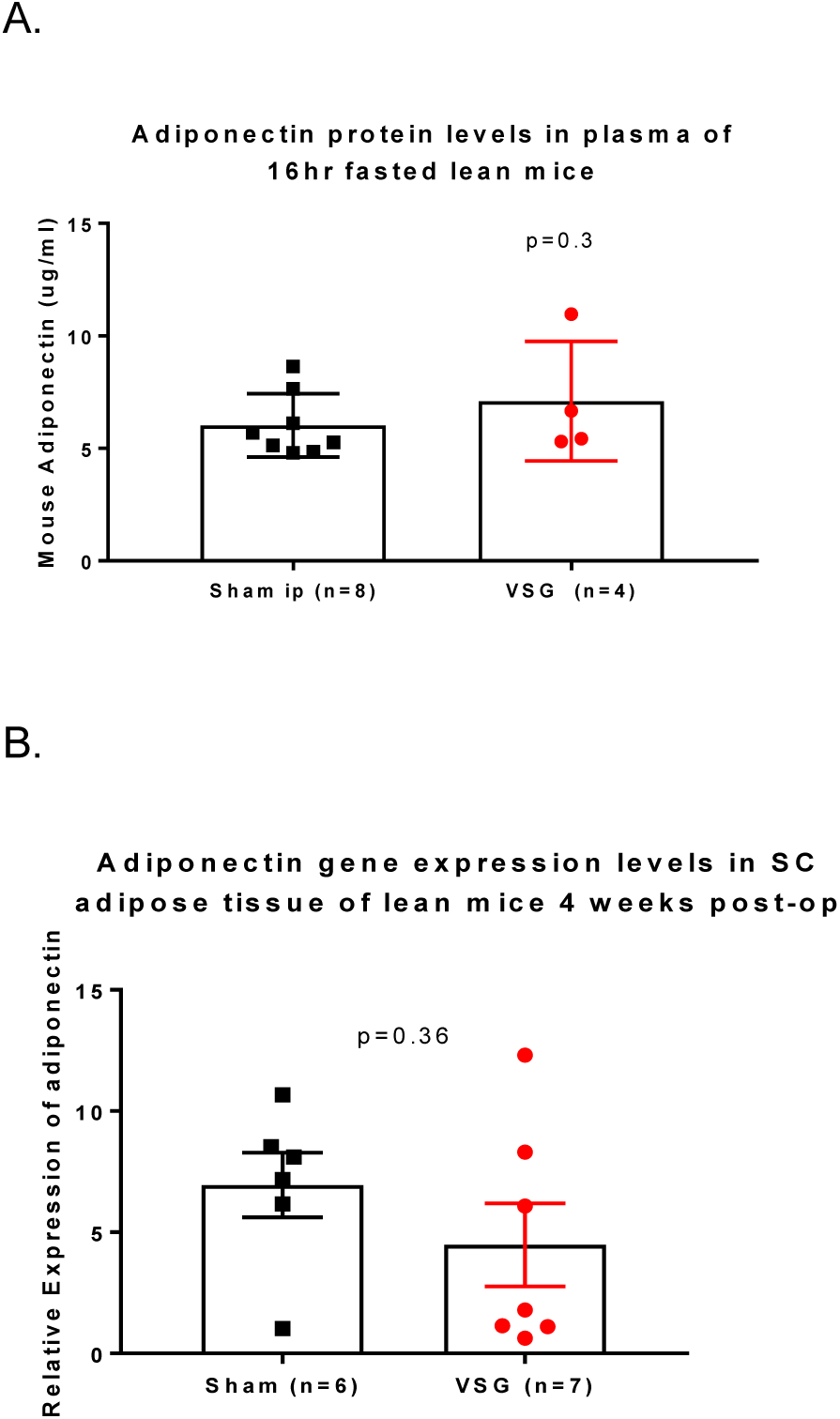
VSG does not increase circulating adiponectin levels (A) Circulating adiponectin was measured in the plasma of 16hr fasted mice, 4 weeks post VSG (B) Quantitative PCR levels of adiponectin gene expression in SC adipose tissue. Data are expressed as means ± SEM

### Hepatocyte nuclear factor 1 alpha (HNF1α) gene expression is not affected by VSG

HNF1α is a transcription factor known to bind to the SGLT2 promoter and to regulate renal proximal tubular function ^36, 37^. We measured HNF1α gene expression in the kidney cortex to detect whether there is a decrease parallel to that observed in SGLT2 gene expression. HNF1α levels were unaffected by VSG (Figure 6).

**Figure 6.**
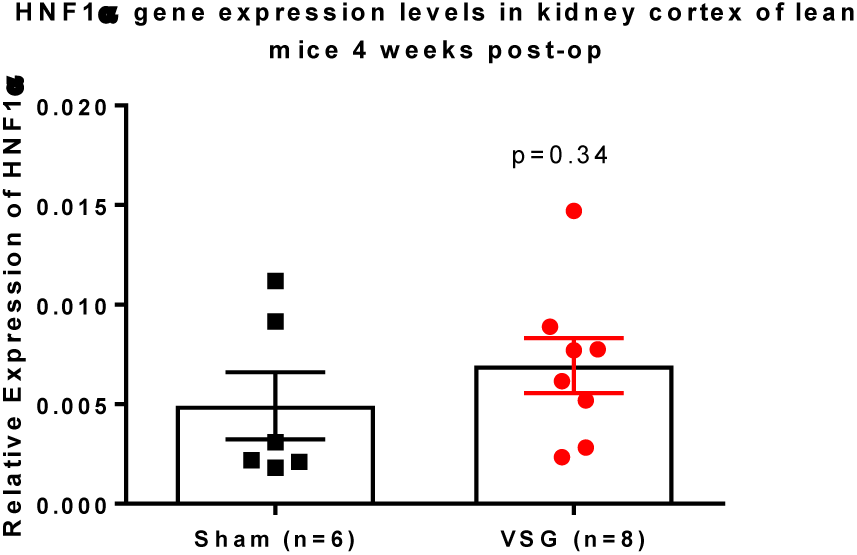
HNF1α transcription factor gene expression is not affected by VSG. Quantitative PCR levels of HNF1α gene expression in kidney cortex. Data are expressed as means ± SEM

## Discussion

This study is the first to our knowledge to demonstrate that the expression of SGLT2 is down regulated in the kidney cortex after bariatric surgery — an inhibition that is preserved in lean animals that had no weight loss at the time of the measurement. Animals that underwent VSG displayed a lowered excursion in glucose and augmented insulin secretion in response to intraperitoneal delivery of glucose, when compared to sham controls. These findings provide important insights into the mechanisms underlying T2D remission following bariatric surgery and point to a potential gut-kidney axis whereby (presently undefined) factors, likely to be released from the intestine, influence gene expression in the kidney.

We originally reported in a meetings proceedings that the stomach sparing duodenal-jejunal bypass in lean rats caused no significant weight loss when compared to sham animals, yet SGLT2 expression was lowered in the kidney cortex^33^. In the present study, we utilised lean mice and performed a different type of bariatric surgery that reduces the size of the stomach rather than rerouting the intestinal tract. Nonetheless, SGLT2 inhibition was preserved across species and surgery, pointing towards communication (which may be direct or indirect) between the gastrointestinal tract and the kidney.

SGLT2 deletion has previously been shown to reduce obesity-associated hyperglycaemia, improve glucose intolerance, and increase glucose-stimulated insulin secretion *in vivo* by preserving β-cell mass via reduction of glucose toxicity, a mechanism that has also been found present during SGLT2 inhibition treatment ^38^. In order to assess if the down-regulated levels of SGLT2 post-VSG result in the same phenotype as SGLT2 inhibitor treatment, we administered dapagliflozin to sham operated mice. The rate of glucose clearance following an intraperitoneal glucose load in the VSG-treated mice remained significantly lower when compared to dapagliflozin-treated sham operated mice. This observation confirms that SGLT2 inhibition likely contributes to bariatric-associated euglycaemia but it is not the sole mechanism involved. Additionally, VSG mice that were also treated with dapagliflozin had identical glucose clearance levels as VSG mice, probably because the treatment has no further additive effect to the already reduced glycaemia levels. A potential explanation for this is that since VSG lowers renal SGLT2 gene expression, and dapagliflozin antagonises the transporter, its action may no longer be observed.

How might VGS lower SGLT2 levels in the kidney? The physiological mechanisms that control SGLT2 expression are not fully understood. There is currently conflicting evidence regarding the role of glucose levels in the glomerulal filtrate of the kidney, although this was initially considered the most prevalent regulator of SGLT2 expression ^39, 40^. However, it was recently reported that hyperglycaemia reduces urinary sodium excretion by enhancing SGLT2 activity ^35^. Adipokines such as tumour necrosis factor-α and interleukin 6 (IL-6) were found to upregulate SGLT2 *in vitro* ^41^, and increased adiponectin was shown to cause activation of PPARδ in the adipose tissue and reduce renal SGLT2 ^35^. Recent evidence showed potential cross-talk between the sympathetic nervous system and SGLT2 regulation 42 while sex and species differences appeared to be influencing SGLT2 expression in rodents ^43^. At the transcriptional level, hepatocyte nuclear factor 1 alpha (HNF1α) is known to bind to the SGLT2 promoter in the kidney, and had been considered a vital transcription factor for SGLT2 expression since HNF1α -null mice have reduced SGLT2 transcript levels in tubular cells ^37^. Moreover, both cyclic AMP and protein kinase A were shown to upregulate SGLT2 at the posttranslational level ^40, 44^.

In order to unravel the causes of the observed SGLT2 inhibition after VSG, we explored a number of different potential mechanisms. The role of glucose was explored by pharmacologically lowering glycaemia in lean mice by injecting them with semaglutide for 7 days. Low glycaemia resulted in reduced SGLT2 expression, yet did not cause the significant inhibition found following VSG. Furthermore, we observed no change in circulating adiponectin, and it is thus unlikely for this to be the mechanism affecting kidney SGLT2 expression. Finally, HNF1α expression levels in the kidney cortex were unaffected by VSG. It is therefore clear that *in vitro* studies in kidney proximal tubular cells are essential to ascribe direct causality of SGLT2 down-regulation and to understand whether this is part of a gut-kidney axis.

The findings described here add to the growing therapeutic attributes of bariatric surgery. We have found that bariatric surgery in lean rodents down-regulates SGLT2 both in gene and protein level, in a weight-independent manner. We also report that exogenous inhibition of SGLT2 action does not fully replicate the positive effects of surgery. More studies are essential to validate whether these outcomes are replicated in diabetic (e.g. obese) rodents, and to distinguish the phenotype between SGLT2 deletion and SGLT2 function inhibition through the use of cell specific knockout mice.

## Acknowledgements

E.A. was supported by a grant from the Rosetrees Trust (M825). G.A.R. is supported by a Wellcome Trust Investigator (212625/Z/18/Z) Award, MRC Programme grants (MR/R022259/1, MR/J0003042/1, MR/L020149/1) and Experimental Challenge Grant (DIVA, MR/L02036X/1), MRC (MR/N00275X/1), and Diabetes UK (BDA/11/0004210, BDA/15/0005275, BDA 16/0005485) grants. This project has received funding from the European Union’s Horizon 2020 research and innovation programme via the Innovative Medicines Initiative 2 Joint Undertaking under grant agreement No 115881 (RHAPSODY) to G.A.R. I.L is supported by a project grant from Diabetes UK (16/0005485).

